# Modular biofabrication of a vascularized skeletal muscle model through endothelialized microvascular seeds

**DOI:** 10.64898/2026.03.31.715476

**Authors:** Fabio Maiullari, Marina Volpi, Nehar Celikkin, Maria Celeste Tirelli, Francesco Nalin, Abhishek Viswanath, Piotr Kasprzycki, Karol Karnowski, Dario Presutti, Wojciech Swieszkowski, Marco Costantini

## Abstract

The clinical translation of engineered skeletal muscle (eSM) for volumetric muscle regeneration is hindered by the challenge of establishing a functional vascular network capable of sustaining its high metabolic demand and ensuring graft survival. Here, we present a bottom-up biofabrication strategy to generate a pre-vascularized in vitro eSM model through the modular assembly of independently matured muscle and vascular compartments. C2C12 myoblasts were encapsulated within core-shell fibers using rotary wet-spinning (RoWS), yielding anisotropically aligned, multinucleated, and contractile myofibers expressing myosin heavy chain and sarcomeric α-actinin. In parallel, gelatin methacryloyl (GelMA)-based microvascular seeds (µVS), pre-endothelialized with human umbilical vein endothelial cells, were engineered to guide rapid and structurally stable vascular formation while preventing uncontrolled capillary self-organization. Fully endothelialized µVS were incorporated into a pro-angiogenic bioink and processed via RoWS to generate tubular vascular fibers with physiological diameters (100–200 μm) and continuous CD31-positive lumens. After independent maturation, muscle and vascular constructs were bioassembled into a hierarchically organized tissue and co-cultured. By decoupling myogenic and angiogenic differentiation, this strategy overcomes medium incompatibility typical of conventional co-cultures, preserving compartment-specific architecture and function and establishing a versatile platform for muscle-vascular modeling and translational muscle repair.

## 1. INTRODUCTION

Skeletal muscle is a highly dynamic and metabolically demanding tissue whose function relies on the tight coupling between aligned myofibers and a dense, hierarchically organized vascular network [1]. Beyond supplying oxygen and nutrients and removing metabolic waste, the vasculature actively regulates muscle homeostasis through reciprocal signaling and cell–cell crosstalk with myogenic cells [2–6]. This bidirectional interaction becomes particularly evident during physiological remodeling, such as exercise-induced angiogenesis and post-injury repair, where coordinated responses from both endothelial and muscle cells are essential for restoring tissue structure and function [3,7,8]. Thus, skeletal muscle function depends not only on its metabolic support but also on the integration of vascular and myogenic compartments within a precisely organized architecture.

In addition to this biological interdependence, skeletal muscle performance is intrinsically linked to its hierarchical structural organization, spanning from sarcomeric arrangement to the alignment of multinucleated myofibers and their assembly into fascicles. This anisotropic architecture underlies efficient force generation, uniaxial contraction, and coordinated load transmission. Accordingly, biomimetic skeletal muscle engineering requires not only cellular alignment but also preservation of multi-scale structural organization together with functional vascular integration. While these features are finely orchestrated in native tissue, reproducing such integrated muscle–vascular complexity in engineered skeletal muscle (eSM) constructs remains a major challenge. The clinical relevance of this challenge is particularly evident in volumetric muscle loss (VML), defined as the loss of more than 20% of the original tissue mass [9]. In this context, the complexity of integrating a spatially organized and functional vascular network within eSM has significantly hindered the development of clinically relevant constructs. Following implantation, eSM vascularization typically depends on host vessel ingrowth through angiogenic sprouting, a process that progresses slowly (approximately 5–17 μm h⁻¹) [10,11]. Consequently, constructs exceeding the diffusion limit are prone to early ischemia and necrosis before sufficient perfusion is established [12]. To address this limitation, pre-vascularization strategies have emerged as a promising approach to accelerate host vessel anastomosis, enhance perfusion, and improve graft survival and functional integration [13,14].

To date, most pre-vascularized eSM models rely on the co-encapsulation of muscle progenitors and endothelial cells within three-dimensional hydrogels or porous scaffolds using biofabrication techniques such as extrusion bioprinting, inkjet printing, or stereolithography [15]. However, the simultaneous culture of these distinct cell populations,each requiring specific biochemical cues, matrix properties, and differentiation timelines,often compromises both myotube maturation and vascular organization [16]. Compartmentalized strategies that enable independent optimization of myogenic and angiogenic microenvironments prior to integration therefore represent a more effective solution. Although previous compartmentalized approaches have demonstrated enhanced regeneration and host vascularization *in vivo* [17], the faithful recreation of aligned, tubular, vessel-like structures running parallel to myofibers, a defining feature of native muscle, remains elusive. As a result, current eSM models are largely confined to millimeter-scale dimensions, limiting their translational potential for volumetric muscle repair.

In our previous work, we developed a rotary wet-spinning (RoWS) platform capable of rapidly producing highly anisotropic muscle constructs exhibiting advanced myogenic differentiation and expression of tissue-specific markers, including myosin heavy chain (MHC), laminin, and titin [18,19]. This scalable approach enabled the fabrication of decimeter-scale histomimetic constructs that demonstrated regenerative potential in a murine tibialis anterior volumetric injury model, promoting graft engraftment and integration with host muscle tissue [20].

Building on this foundation, the present study aims to further enhance the functional and translational relevance of eSM by incorporating vascular features that recapitulate native muscle–vasculature organization. We engineered stable, hollow, vessel-like microstructures with physiologically relevant diameters (100–200 μm) using gelatin methacryloyl (GelMA)-based microvascular seeds (µVS) produced via a microfluidic step emulsification process. Following endothelialization with a continuous monolayer of human umbilical vein endothelial cells (HUVECs), the µVS were embedded within a pro-angiogenic bioink and processed using the RoWS platform to promote maturation of lumenized microvessels. In parallel, skeletal muscle modules were independently fabricated by encapsulating C2C12 myoblasts within core–shell fibers and cultured under myogenic conditions to generate aligned, multinucleated, contractile myofibers. After independent maturation, vascular and muscular components were assembled into a unified, anisotropically organized construct. This modular biofabrication strategy preserves tissue-specific structure and function while recapitulating the hierarchical architecture of native vascularized skeletal muscle, providing both a translational platform for volumetric muscle regeneration and a versatile *in vitro* model to study muscle–endothelium crosstalk.

## 2. MATERIALS AND METHODS

### 2.1 3D RoWS setup

Core/shell hydrogel fibers were fabricated and 3D spatially deposited using a custom-built 3D wet-spinning bioprinter developed in our laboratory and previously employed and described in earlier publications [1–4]. Briefly, the whole platform consists of: i) an extrusion system composed of a Microfluidic Printing Head (MPH), bearing a crosslinking bath microtank with a co-axial nozzle placed at the bottom of it for the immediate gelation of core/shell fiber, ii) a rotating drum collector (diameter = 25 mm, length = 180 mm), and iii) an X-axis (travel range = 160 mm) for the sequential extrusion process. The entire system was controlled by an Arduino Mega board and custom Python software. The MPH was fabricated using an Elegoo Mars 5 Ultra 3D printer and BioMed Clear Resin (Formlabs, FLBMCL01). The internal microchannels within the MPH base were printed with an 800 µm internal diameter, while the coaxial outlet incorporated 600 µm-diameter channels for both the core and shell flow paths. Finally, the MPH was equipped with a crosslinking bath microtank and mounted on the bioprinter’s X-axis arm. The inlets of the printing head were connected to programmable microfluidic syringe pumps (neMESYS low pressure, Cetoni GmbH, Germany) via autoclavable Teflon tubing (ID = 800 μm).

### 2.2 Cell culture

C2C12 murine myoblasts and human umbilical vein endothelial cells (HUVECs) were cultured under standard conditions at 37 °C in a humidified atmosphere containing 5% CO₂. C2C12 cells were expanded in high-glucose Dulbecco’s Modified Eagle Medium (DMEM) supplemented with 10% fetal bovine serum (FBS) and 1% penicillin-streptomycin (Pen/Strep), whereas HUVECs were maintained in complete endothelial growth medium supplemented with growth factors (EASY Endothelial Cell Growth Medium kit, PeloBiotech). Both cell types were passaged upon reaching approximately 60% confluency using 0.25% Trypsin-EDTA (Gibco), and replated in new culture dishes for expansion. Cells were expanded until the quantities required for the formulation of the respective bioinks were reached.

### 2.3 GelMA synthesis

GelMA was synthesized following a previously published protocol [5]. Briefly, gelatin (type A3, ∼300 Bloom, derived from porcine skin) was dissolved in PBS at a concentration of 10% (w/v) at 60°C. Methacrylic anhydride (MA) was then added dropwise at a ratio of 0.08 mL per gram of gelatin under vigorous stirring. The reaction was allowed to react for 2 hours. The resulting solution was subsequently diluted five-fold with warm PBS and dialyzed using 12–14 kDa molecular weight cutoff dialysis tubing (Spectrum Laboratories) for 6 days at 50 °C to remove residual MA and by-products. After dialysis, the GelMA solution was lyophilized and stored at −20 °C until further use.

### 2.4 RoWS of engineered skeletal muscle

Core/shell hydrogel fibers were fabricated using the custom co-axial MPH. The core bioink consisted of 14 ×10^6^ C2C12 myoblasts/mL suspended in a sterile-filtered solution of 1.4% w/v fibrinogen from bovine plasma in 25 mM HEPES buffer containing 150 mM NaCl. The shell biomaterial ink was formulated by dissolving 2% w/v LMW-ALG, 0.8% HMW-ALG-RGD (ALG-RGD) in 25 mM HEPES. The alginate RGD reaction was performed as previously described [6]. The inks were supplied to the MPH nozzle at controlled flow rates to establish a stable co-flow within the extrusion nozzle while the crosslinking bath microtank was pre-filled with 0.6 M CaCl_2_ solution. For all cellular experiments, the core and shell flow rates were maintained at Qc = 160 µl min^−1^, Qs =320 µl min^−1^ (Qc = flow rate core, Qs = low rate shell), respectively. Upon extrusion, the core/shell fibers underwent instantaneous gelation upon contact with the CaCl₂ solution at the nozzle tip. The resulting core-shell hydrogel fiber was initially pulled gently upwards using a tweezer until reaching the surface of a rotating Teflon drum. Continuous extrusion of core-shell fibers combined with rotational collection enabled the formation of anisotropic bundles. Upon contact with the drum, a hydrogel fiber is continuously extruded from the nozzle and collected onto the drum, forming a bundle. The drum rotation speed was set at 60 rpm, and the fiber number in each bundle was kept constant (30 threads in each bundle). After extrusion, the samples were collected from the rotating drum and subjected to secondary crosslinking of the core bioink by incubation in a 2 U/mL thrombin in 25 mM HEPES buffer solution for 30 min at 37 ^◦^C. The bundles were cultured at 37 °C in a humidified atmosphere with 5% CO₂. For the first 4 days, constructs were maintained in high-glucose DMEM supplemented with 20% FBS, 1% Pen/Strep, 1% L-Glutamine and 0.5 mg/mL ε-aminocaproic acid (ACA). At the cell confluence stage, the medium was switched to high-glucose DMEM containing 2% horse serum, 1% Pen/Strep, GlutaMAX and 0.5 mg/mL ACA for the following days of culture.

### 2.5 High-throughput microvascular seed (µVS) production

#### Millipede chip production

The masters for the microfluidic chips were fabricated by micropatterning a silicon wafer using standard two-layer photolithography [7]. First, a thin layer was obtained by pouring a few mL of SU8-2050 on a silicon wafer to produce thin microchannels (spin-coating at 4000 rpm for 30 s). The resist was soft-baked at 65 °C for 2 minutes and 95 °C for 6 minutes, and then exposed to UV light through a high-precision photolithography mask. The patterned master was cured at 65 °C for 1 minute, then at 95 °C for 6 minutes, and finally immersed in SU8 developer for 5 minutes to remove the unexposed resist. A similar procedure was followed to create the thicker layer, this time, coating the wafer with SU8-2150 (spin-coating at 2500 rpm for 30 s). The master was baked for 7 minutes at 65 °C and 45 minutes at 95 °C. Alignment markers on the second lithography mask were used to align the features of the first layer with the ones of the second layer. The resist was exposed to UV light through the mask, and then cured at 65 °C for 5 minutes and 95 °C for 15 minutes. The unexposed resist was removed in a bath of SU8 developer for 15 minutes and then the master was thermally treated at 195 °C for 30 minutes. The resulting master presented microstructures with a thickness of 37 µm for the thin layer and a 250 µm for the thick layer. The microfluidic chips were prepared by pouring PDMS (10:1 with a crosslinking agent) onto the patterned surface of the master and baking the PDMS for 2 hours at 75 °C. Later, the cured PDMS was removed from the master and sliced along the chip outline. Inlets and outlets of the chip were obtained using a biopsy puncher, creating holes with a diameter d = 1 mm. The PDMS was bonded to clean glass slides through a standard plasma bonding [8,9]. The hydrophobic surface modification of the chip was obtained by injecting a 5% v/v solution of Silane (Trichloro(1H,1H,2H,2H-perfluorooctyl) silane, ref. 448931, Sigma-Aldrich) in HFE-7500 (3M, USA). After 20-30 minutes,the Silane solution was removed from the channels with a stream of compressed air.

#### Beads production

Two glass Hamilton syringes were prepared for water-in-oil droplet generation. Syringe_1 was filled with the disperse phase consisting of 5% w/v GelMA supplemented with 0.01% w/v Irgacure 2959 photoinitiator in PBS, and the Syringe _2 was filled with the continuous phase composed of Novec 7500 (3M) containing 1% w/w PFPE–PEG–PFPE surfactant [10,11]. Each syringe was connected to a 25G stainless steel needle and linked to the millipede chip inlet via Teflon tubing (0.8 mm O.D., 0.5 mm inner diameter). Droplet formation was controlled using precision syringe pumps (Cetoni Nemesys S, Germany) to set the flow rate of both the dispersed and continuous phases to 100 µL/min. The resulting hydrogel beads were collected in 200 µL aliquots into 1.5 mL microcentrifuge tubes and subsequently crosslinked by exposure to UV light (365 nm, 5 mW/cm²) for 15 minutes.

#### Beads purification

Following UV crosslinking, the beads were washed with Novec 7500 containing 20% v/v 1H,1H,2H,2H-perfluorooctanol (PFO; Thermo Scientific) for 20 seconds to destabilize residual emulsion. Excess fluid was removed from the bottom of the tube, and the bead pellet was sequentially washed four times with pure Novec 7500 and four times with sterile PBS, using centrifugation steps at 200 × g for 5 minutes between each wash. After the final wash, beads were resuspended in a small volume of PBS to obtain a maximally packed bead suspension and subjected to UV sterilization for 2 hours under a laminar hood.

#### Cell seeding on the beads

To generate the final microvascular seeds, HUVECs were homogeneously seeded onto the surface of the GelMA beads. Specifically, 100 µL of beads (50 000 beads) were mixed with 2 mL of a cell suspension to achieve a final concentration of 2 × 10⁶ cells/mL, and incubated under gentle rotation at 37 °C for 4 hours to promote uniform cell attachment.

### 2.6 RoWS of microvessels

Engineered vascular fibers were fabricated using the same core–shell wet-spinning protocol described in section 2.4 ‘RoWS of engineered skeletal muscle’. The formulation of the shell bioink and all spinning parameters, including extruder configuration, flow rates, rotational speed, and crosslinking conditions, were maintained unchanged.

The core bioink composition was specifically optimized to support endothelial cell attachment and vascular morphogenesis. After cell seeding, microvascular cells were pelleted by centrifugation and gently resuspended in 400 µL of endothelial growth medium. The obtained microvascular seed suspension was then mixed with a stock fibrinogen solution containing a single-cell HUVEC suspension. The resulting mixture was formulated to achieve final concentrations of 7 mg/mL fibrinogen and 10^7^ cells/mL. During the spinning process, fibers were collected on the rotating drum, then transferred into a 6-well plate and subjected to enzymatic crosslinking of the fibrinogen-based core bioink. This was performed by incubating the constructs in a thrombin solution (2 U/mL in 25 mM HEPES buffer) for 30 minutes at 37 °C. The engineered vascular bundles were subsequently cultured at 37 °C in a humidified atmosphere with 5% CO₂ in complete Endothelial Cell Growth Medium for up to 12 days.

### 2.7 Vascularized Skeletal Muscle Assembly

To generate an integrated biomimetic construct composed of aligned muscle tissue integrated with a microvascular network, engineered muscle and vascular bundles were combined and structurally integrated into a single composite bundle. Specifically, on day 18 of skeletal muscle maturation and day 12 of vascular bundle culture, the respective engineered tissues, muscle fibers, and tubular microvessels, were manually co-positioned to ensure direct contact and spatial continuity between compartments. 5% w/v GelMA solution, previously supplemented with 0.05% w/v Irgacure 2959 in HEPES, was gently applied to embed and stabilize the interface between the two tissue types. The assembly was then exposed to UV light (365 nm, 12 mW/cm²) for 1 min to induce photopolymerization of the GelMA. Following crosslinking, the integrated constructs were maintained in a co-culture medium consisting of endothelial growth medium, with FBS replaced by 2% horse serum (EASY Endothelial Cell Growth Medium kit, PeloBiotech), supplemented with 0.5 mg/mL ACA. This condition supported the simultaneous culture of both skeletal muscle and endothelial compartments, ensuring cohesive integration while preserving their individual architecture and cellular organization.

### 2.8 Optical coherence microscopy (OCM)

Morphological characterization of 3D spun samples was performed using high-resolution OCM [12]. A custom-designed spectral OCM system used a commercial femtosecond laser (Fusion Femtolasers, Austria) with a 130 nm full bandwidth (resulting in an axial resolution of 2.2 μm) and a 795 nm central wavelength as a light source. The Michelson interferometer configuration employed a fiber-based beam splitter with a 75/25 splitting ratio. The light reflected from the sample was combined with light reflected from the mirror in the reference arm, and the resulting interference signal was recorded with a high-speed 2048-pixel line-scan camera spectrometer (Cobra-S 800, Wasatch Photonics, USA). A 4× microscope objective (Olympus Plan Fluorite, NA = 0.13, WD = 17.3 mm) was used for sample illumination. The pupil was exposed to a 6 mm (1/e2) Gaussian beam, leading to a lateral resolution of 4.5 μm, and a confocal parameter f ≈ 100 μm. The sample was illuminated with an average optical power of 0.5 mW. Lateral laser beam scanning was performed using a pair of galvoscanners (Thorlabs Inc., USA). 3D volumes typically extending over an area of 1.0 mm × 1.0 mm × 1 mm were acquired with a pixel density of 1024 × 1024 × 2048 pixels for X-, Y-, and Z-axes, respectively. The imaging depth, limited by light penetration and multiple scattering, was evaluated to be 800 μm. Specific stand-alone software, designed in LabVIEW, enabled the monitoring of a scanning protocol, the coordination of the camera and scanners, and data processing, including steps such as spectrometer calibration, numerical dispersion compensation, and spectral shaping. Data analysis enabled the volumetric reconstruction of the fine structure of the investigated spun fiber bundles.

### 2.9 Scanning Electron Microscopy (SEM)

The cross-sectional architecture of the core-shell fibers was analyzed by scanning electron microscopy (SEM) using a Nova Nano SEM system (FEI Company, USA). Prior to SEM analysis, samples were fixed overnight in 4% paraformaldehyde (PFA) and dehydrated using a graded ethanol series (50–100%), with 1 h incubation at each step. Dehydrated samples were then subjected to supercritical CO₂ drying using a supercritical point dryer (AutoSamdri-815, Tousimis, USA) to prevent collapse of the fibrinogen-based core and preserve the native cross-sectional morphology. Finally, samples were mounted on aluminum stubs using double-sided conductive carbon tape and sputter-coated with a 7 nm-thick gold layer to minimize surface charging. SEM imaging was performed at an accelerating voltage of 5 kV.

### 2.10 Rheology

The rheological properties of the cell-free prepolymer hydrogel formulations used for bioink preparation were analyzedusing a Kinexus Pro rotational rheometer (Malvern Panalytical Ltd.) equipped with a cup-shaped outer cylinder and rotating inner cylinder, in which the diameter of the outer cylinder is 27.5 mm and the diameter of the inner cylinder is 25 mm. The rheological test was performed at room temperature (25 °C). The shear viscosity and shear stress measurements for the shell, muscle, and microvascular ink formulations were performed in shear rate-controlled mode over a shear rate range of 0.1–1000 s⁻¹.

### 2.11 Mechanical characterization

Compressive tests on bulk hydrogels were performed after 24 h of incubation in 25 mM HEPES buffer (n = 4 for each ink formulation) using a Dynamic Mechanical Analyzer (DMA Q800, TA Instruments). Hydrogels were prepared as cylindrical specimens (8 mm diameter × 6 mm height; 300 µL volume) prior to testing. Measurements were carried out at room temperature with an initial preload of 0.001 N. A compressive force ramp of 0.01 N·min⁻¹ was applied until a maximum strain of 20% was reached. The elastic modulus (E) was determined from the stress–strain (σ–ε) curves as the slope of the linear region between 0 and 5% strain (R² > 0.9). The maximum stress (σ₂₀%) was defined as the stress value at 20% strain.

### 2.12 Immunofluorescence

On specified time points (skeletal muscle bundles: day 10 and day 18, vascular bundles: day 6 and day 12, assembled bundles: day 7), core–shell fiber bundles were fixed in 4% (w/v) paraformaldehyde (PFA) overnight at 4 °C for immunofluorescence analysis, thoroughly washed, and stored in 25 mM HEPES buffer at 4 °C until staining. Samples were permeabilized immediately before staining with 0.3% (v/v) Triton X-100 in PBS for 1 hour at room temperature and immersed in blocking buffer (5% w/v bovine serum albumin, BSA, in PBS) for 2 h at RT to reduce non-specific antibody binding. The samples were incubated overnight at 4 °C with primary antibodies diluted in 0.5% BSA in PBS. The following primary antibodies were used: mouse anti-CD31 Alexa Fluor 647-conjugated (1:50; Abcam, ab28364), sheep anti-Von Willebrand Factor (1:100, 11713, Abcam) for endothelial cells; mouse anti-Myosin (Skeletal) (1:2; DSHB, Hybridoma Product MF20), rabbit anti-Laminin antibody (1:60, Sigma-Aldrich – L9393) and mouse anti-Sarcomeric Alpha Actinin antibody (1:25, ThermoFisher Scientific – MA1-22863) for skeletal muscle fibers. F-actin was detected using Alexa Fluor 488-conjugated phalloidin (1:40, Thermo Fisher Scientific, A12379). The samples were then washed three times with PBS (10 min each) and incubated for 4 h at room temperature with the corresponding Alexa Fluor secondary antibodies (488, 568, or 647; 1:1000, Thermo Fisher Scientific). Cell nuclei were counterstained with DAPI (1:1000) for 30 minutes at room temperature, followed by final washes in PBS. Immunostaining of µVS was performed using a standard immunofluorescence protocol for 2D cell cultures [31]. Briefly, the µVS were fixed in 4% (w/v) PFA for 15 min at room temperature, permeabilized with 0.1% (v/v) Triton X-100 in PBS for 15 min, blocked with 5% (w/v) BSA in PBS for 20 min and subsequently incubated with Alexa Fluor 488-conjugated phalloidin (1:40; Thermo Fisher Scientific, A12379) for 2 h at room temperature. Nuclei were counterstained with DAPI (1:1000) for 10 min at room temperature before imaging. Fluorescent imaging was performed using a confocal laser scanning microscope (Nikon A1R).

### 2.13 Live/dead assay

Following the assembly phase, the engineered skeletal muscle and vascular constructs were further cultured for one week under different conditions to evaluate cross-compatibility between the two compartments. Specifically, muscle (day 18) and vascular 3D constructs (day 12) were maintained either in their respective growth media or in a co-culture medium (see Section 2.7 for details). At end of culture, cell viability was assessed using the LIVE/DEAD Cell Imaging Kit (Thermo Fisher Scientific, R37601) according to the manufacturer’s protocol. Briefly, an equal volume of the 2× working solution, prepared by mixing components A (Live Green) and B (Dead Red), was added directly to the samples and incubated for 15 min at room temperature (20–25 °C) before confocal imaging.

### 2.14 Image processing

Image processing and quantitative analyses were performed using ImageJ (NIH, Bethesda, MD, USA). The thickness of myotubes, as well as the spatial distribution of the core and shell components, and the sarcomeric length (i.e. inter Z-disk spacing), were quantified using the Plot Profile tool. Quantitative evaluation of GelMA microbeads diameter and the lumen size of vascular structures were determined using the Measure function.

### 2.15 Spontaneous contraction analysis

Videos of spontaneous contractions were recorded using the Recording function of NIS-Elements software (Nikon) and saved in .avi format for subsequent analysis. Contractility characterization was performed using the Myocyter v1.3 plugin for ImageJ, according to the developer’s user manual [13]. A preliminary Pretest (for size and threshold) was executed to optimize the recognition of contracting regions and to exclude the background. During this step, the appropriate size range and intensity threshold were adjusted to ensure that only the turquoise-highlighted areas corresponding to the contracting portions of the sample were selected, while non-contractile background regions were completely excluded. Following the pretest optimization, the Evaluation function was run using the identified size and threshold parameters to extract quantitative contractility data. The results were automatically compiled and exported as Excel files for subsequent analysis

### 2.16 Gene Expression Analysis

At the designated experimental time points, the spun samples were transferred into sterile 2 mL RNase-free tubes, washed in PBS for 5 min, and digested in alginate lyase (150 µg/mL) for 15 min at 37 °C. Samples were then centrifuged for 5 min at 300 × g, rapidly frozen in liquid nitrogen, and stored until RNA extraction. Total RNA was extracted from samples of each experimental condition using a mortar and pestle and liquid nitrogen. The pulverized material was collected with 700 µL of TRIzol reagent (Invitrogen, Life Technologies) and transferred into a new RNase-free tube. It was then purified using the RNeasy Mini Kit (Qiagen, 74106) according to the manufacturer’s protocol. RNA concentration and purity were assessed using a NanoDrop UV-Vis spectrophotometer (Thermo Fisher Scientific). For reverse transcription, 500 ng of total RNA were converted into cDNA using the High-Capacity cDNA Reverse Transcription Kit (Applied Biosystems - 374966) in a total reaction volume of 20 µL. The resulting cDNA was then diluted to a final volume of 50 µL. Quantitative real-time PCR (qRT-PCR) was performed using the RT-PCR Mix SYBR (A&A Biotechnology-2008-1000) on a LightCycler 96 system (Roche). All reactions were carried out in duplicate in a total volume of 25 µL. Gene expression was evaluated using the ΔΔCt method. Fold change values were subsequently expressed relative to the control group by subtracting 1 from the 2^−ΔΔCt values, so that positive and negative values indicate upregulation or downregulation relative to baseline, respectively. TBP was used as a housekeeping gene for muscle samples and GUS for microvascular samples. Primer sequences are reported in Supplementary Table 1.

### 2.17 Statistical analysis

Statistical analyses were performed using GraphPad Prism (GraphPad Software, La Jolla, CA, USA). Data are presented as mean ± standard deviation (SD). For qRT-PCR analyses, differences among multiple time points were evaluated using one-way ANOVA. Comparisons between individual experimental groups for myotube thickness measurements were performed using Student’s t-test. A p-value < 0.05 was considered statistically significant.

## 3. RESULTS AND DISCUSSION

### 3.1. RoWS of highly aligned core-shell hydrogel fibers

The workflow for generating a vascularized skeletal muscle model relies on two independent RoWS biofabrication processes, enabling the separate generation of muscle and vascular-like compartments before their integration into a single tissue construct. Skeletal muscle fibers were fabricated using a myogenic bioink and cultured in a muscle-specific medium to induce myoblast differentiation. In parallel, GelMA microbeads pre-coated with endothelial cells were embedded in an angiogenic bioink and cultured for 12 days, enabling the formation of lumenized microvascular networks with a continuous endothelial monolayer at the fiber core-shell interface.

Upon reaching their respective maturation endpoints, pre-formed muscle and vascular modules were assembled into a spatially defined construct to create a vascularized skeletal muscle model. Core-shell fibers were fabricated using the rotary wet-spinning (RoWS) bioprinter developed by our group. The platform was equipped with a newly designed microfluidic co-axial extrusion device (**Figure 1a**). The printing head was fabricated as a single, monolithic unit using a biocompatible, USP Class VI-certified, resin that offers high precision, chemical resistance, and compatibility with standard sterilization methods. This MPH design ensured structural robustness and eliminated the need for manual assembly and nozzle alignment steps, thanks to a single-step DLP 3D-printing fabrication process that maintained the intrinsic coaxial configuration of the internal channels [4]. The device features independent inlets for the core and shell phases, enabling precise and stable core-shell spatial patterning of the bioinks during extrusion. By modulating bioprinting parameters, such as the ink flow rate or the collection drum rotational speed, the dimensions of the core and shell phases can be independently and precisely tuned. A thorough characterization of the fiber extrusion process has been published in an article [4]. The core phase was formulated with a fibrinogen-based bioink, selected for its biocompatibility and ability to provide a cell-supportive microenvironment. The shell phase consisted of an alginate-based hydrogel optimized to provide mechanical stability and controlled diffusion. A representative image of the resulting core-shell bundle after thrombin-mediated crosslinking of the fibrinogen-based core is presented in **Figure 1b**. The pre-hydrogel formulations exhibited distinct rheological characteristics, with alginate showing significantly higher shear viscosity than the fibrinogen-based core inks used for muscle and microvascular constructs. Additionally, as shown in **Figure 1c**, shell ink exhibited a constant shear viscosity in the studied range while the core inks (muscle and microvascular inks) revealed a mild shear thinning behavior. Upon simultaneous extrusion of alginate and fibrinogen into a calcium chloride (CaCl₂) crosslinking bath, the shell phase undergoes instantaneous ionic gelation, achieving rapid structural stabilization and mechanical robustness to efficiently encapsulate the fibrinogen in a core-shell configuration. The extruded hydrogel filament is then continuously collected onto the rotating drum to form parallel, aligned bundles. The optimized flow rate ratio (Q_core_ = ½ Q_shell_) facilitated uniform bioink distribution, resulting in fibers with a controlled diameter [1]. The influence of flow rate on fiber dimensions was evaluated by comparing constructs produced under low (Q_core_ = 80 µL/min, Q_shell_= 160 µL/min) and high (Q_core_ = 160 µL/min, Q_shell_ = 320 µL/min) flow rates. At low flow rates, fibers had an average outer diameter of approximately 280 µm and a core diameter of ∼180 µm.

**Figure 1.**
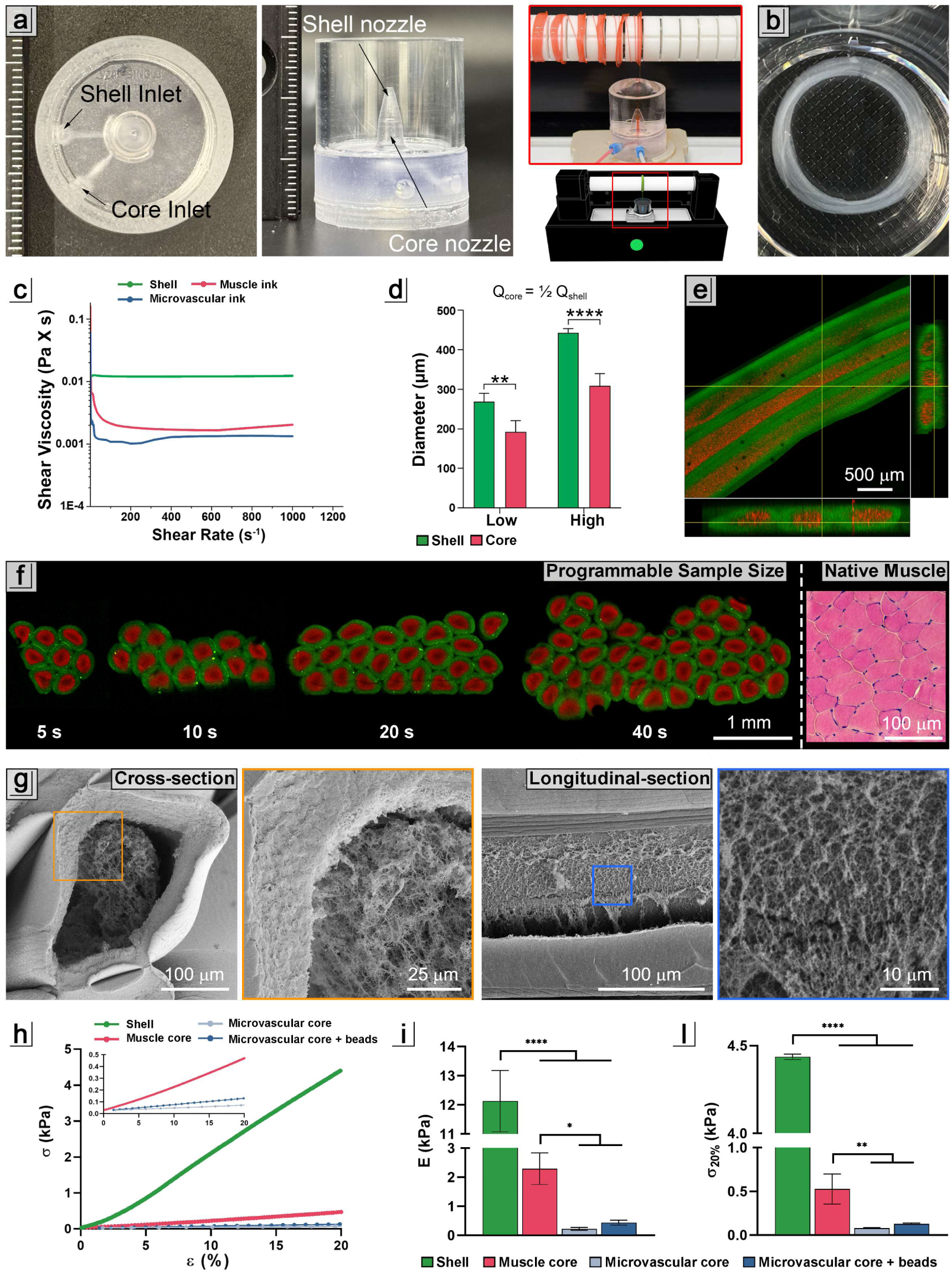
3D rotary wet-spinning (RoWS). a) Representative images of the custom 3D-printed extrusion head, shown from top and side views. A schematic rendering of the rotary wet-spinning printer is shown, along with a representative image acquired during the spinning process, in which core-shell fibers (red) are collected onto the rotating drum. b) Representative image of a core-shell fiber bundle after thrombin-mediated crosslinking of the fibrinogen-based core. c) Viscosity profiles of the shell, muscle and microvascular inks measured as a function of shear rate (1–1000 s⁻¹). The shell ink (green), composed of 2% low molecular weight alginate and 0.8% high molecular weight RGD-alginate, the muscle bioink (red), containing 14 mg/mL fibrinogen and 0.2% low molecular weight alginate, and the microvascular bioink (blue), containing 7 mg/mL fibrinogen; d) Quantification of core diameter and shell thickness at low and high flow rates of the shell (Q_shell_) and core (Q_core_); e) Fluorescence images of a core-shell hydrogel bundle, showing a continuous and homogeneous core matrix (red) surrounded by a stable shell phase (green) with a well-defined boundary. Cross-sectional views confirm effective compartmentalization of core and shell; f) Confocal fluorescence images of cross-sections from bundles composed of increasing numbers of collected fibers, illustrating the tunability of sample size as a function of fiber number. Extrusion times corresponding to each condition (5, 10, 20, and 40 s; drum speed 60 rpm) are indicated, demonstrating temporal control over fiber deposition. A representative cross-section of native skeletal muscle is shown for comparison; g) Scanning electron microscopy (SEM) images of core-shell fibers acquired in cross-section and longitudinal section, highlighting the distinct internal microstructures of the alginate-based shell and the fibrinogen-based core, as well as the longitudinal organization of the fiber; h–l) Mechanical characterization of bulk constructs prepared using the same formulations as the shell ink, muscle core ink, and microvascular core ink, with and without beads. Representative stress-strain (σ-ε) curves (h), elastic modulus (E, kPa) (i), and maximum stress at 20% strain (σ₂₀%, kPa) (l) are shown. *p < 0.05, **p < 0.01, ***p < 0.001, ****p < 0.0001.

In contrast, high flow rates yielded fibers with an average outer diameter of approximately 450 µm and a core diameter of ∼300 µm. These findings confirmed a direct correlation between total flow rate and fiber diameter, with increased flow rates yielding proportionally larger fibers (**Figure 1d**). Precise control over shell hydrogel distribution yielded a uniform outer layer, thereby contributing to the structural integrity and long-term stability of the fibers, as demonstrated by confocal microscopy (**Figure 1e**). Overall, this setup enables reproducible, continuous generation of anisotropic constructs with a well-defined core-shell architecture and consistent material compartmentalization.

A key advantage of the RoWS approach is the ability to readily program the final sample size by controlling the extrusion time for each sample. As shown in **Figure 1f**, constructs ranging from a limited number of aligned fibers to densely packed bundles can be generated in a highly reproducible manner. Given the constant rotational speed of the collection drum (60 rpm), the number of deposited fibers scales proportionally with extrusion time, enabling temporal control over construct size. Notably, the fiber is extruded continuously, introducing a minor variability in fiber number (±1–3) per construct during the translational movement of the extrusion head between consecutive samples. This does not affect the structural organization or overall construct quality, but it represents a parameter that could be further optimized in future platform refinements. Confocal cross-sectional images (shell in green, core in red) demonstrate that increasing fiber number directly translates into larger tissue-like assemblies while preserving structural alignment and compartmentalization. Notably, the resulting cross-sectional organization closely resembles the architecture of native skeletal muscle, as highlighted by comparison with hematoxylin and eosin (H&E) staining of native muscle tissue, underscoring the biomimetic potential of the system. Scanning electron microscopy (SEM) further confirmed the structural integrity and material-specific organization of the core-shell fibers (**Figure 1g**). Cross-sectional and longitudinal images revealed a clear spatial segregation between the alginate-based shell and the fibrinogen-based core. The shell exhibited a compact, low-porosity morphology, providing mechanical containment and structural definition, whereas the fibrinogen core displayed a highly fibrillar, porous microarchitecture. This distinct internal organization supports efficient encapsulation while providing a permissive microenvironment that may enhance cellular viability and growth within the core compartment. To investigate the mechanical properties of the proposed bioink formulations (shell, muscle-mimicking core, vessel-mimicking core, and vessel-mimicking core containing GelMA-based beads), confined compression tests were performed on the corresponding crosslinked bulk hydrogels. The resulting stress-strain curves revealed distinct mechanical behaviors between shell and core formulations (**Figure 1h**). Fibrin hydrogels used for the muscle-mimicking bioinks exhibited a compressive elastic modulus of 2.29 ± 0.44 kPa (**Figure 1i**). This value falls within the range reported for hydrogel support matrices used in engineered-driven skeletal muscle regeneration [14]. Similarly, vessel-mimicking bioinks displayed stiffness values (∼0.3 kPa) comparable to those of soft engineered extracellular matrices previously shown to support capillary-like network formation [15]. Alginate-based constructs showed a significantly higher compressive elastic modulus (12.13 ± 0.85 kPa), corresponding to approximately 6-fold and 60-fold increases compared to the muscle- and vessel-mimicking core inks, respectively. These enhanced mechanical properties are consistent with the role of alginate bioinks as supporting materials for the fabrication of wet-spun core–shell microfibers. No significant differences were observed between the vessel-mimicking core ink and the vessel-mimicking core containing GelMA-based beads, indicating that bead incorporation did not alter the intrinsic mechanical properties of the pristine 7 mg mL⁻¹ formulation nor affect crosslinking efficiency. Notably, the vessel-mimicking core with beads exhibited a low standard deviation, suggesting a homogeneous distribution of microbeads within the hydrogel matrix and preserving sample-to-sample reproducibility. Consistently, the maximum stress measured at 20% compressive deformation followed the same trend observed for the elastic modulus, with shell inks reaching higher stress values (∼4 kPa) compared to core formulations (∼0.5-0.1 kPa) (**Figure 1l**). This difference indicates a greater load-bearing capacity of the shell bioinks, while the lower stresses registered for the core inks are consistent with their compliant mechanical profile.

### 3.2 Biofabrication and maturation of anisotropically oriented eSM

To experimentally validate our biofabrication strategy and assess the structural and functional maturation of engineered skeletal muscle (eSM), we first generated anisotropically oriented myofibers using the RoWS platform and monitored their development over time. The timeline in **Figure 2a** illustrates the sequential stages of myogenic differentiation within the biofabricated wet-spun core-shell microfibers. Starting from initial cell encapsulation (T0), myotubes were detectable by day 7 (T7) and progressively matured into a highly aligned, structurally consolidated muscle architecture from day 10 (T10) onwards, further strengthening beyond day 14 (>T14). Brightfield microscopy images (**Figure 2b**) showed a progressive transition from single myogenic precursors (Day 0) to highly aligned, densely packed myotubes (Day 18). Quantitative directionality analysis confirmed this trend, with a marked increase in anisotropy over time. In skeletal muscle constructs, such an increase in directionality can be directly linked to an increase in cell orientation, a crucial feature that ultimately ensures proper functionality and force generation of the engineered myo-constructs [16]. At early time points (Day 0 and Day 1), no detectable orientation was observed, as cells were still in the initial spreading phase and did not yet perceive the anisotropic cues provided by the fiber architecture. As confluence was approached, an emerging alignment became evident by day 4, further strengthened by day 7. This progressive orientation culminated at later stages: at days 10 and 18, the directionality histogram shows a sharp peak at 0°, indicating a predominant uniaxial alignment, a hallmark of native skeletal muscle organization (**Figure 2b**). Immunofluorescence staining for myosin heavy chain (MHC) and laminin (**Figure 2c**) confirmed the progressive myogenic differentiation of the constructs. By day 10, multinucleated myotubes were already detectable, exhibiting MHC-positive fibers and basement membrane deposition, as evidenced by laminin staining. After 18 days of culture, myotubes appeared more compact and densely arranged in the fiber, with continued expression of MHC and laminin along their length, indicative of ongoing maturation and SM tissue consolidation [17]. In line with the observed structural maturation, quantitative analysis of myotube thickness revealed a significant increase at later differentiation time points (**Figure 2d**). At day 10, myotubes exhibited an average diameter of approximately 15 µm, whereas by day 18 the diameter distribution shifted toward ∼18 µm. Although this increase may appear modest, it corresponds to an approximate 40% increase in myotube volume, indicating continued fiber growth and stabilization within the core. Distribution analysis further supported this trend: whereas most myotubes at day 10 clustered around 15 µm, day 18 constructs exhibited a reduced proportion of thinner fibers and a greater representation of thicker myotubes (25 µm), along with the appearance of larger structures exceeding 35 µm. While the changes in relative frequency are moderate, the shift and redistribution toward higher-caliber myofibers, along with the emergence of larger myotubes, collectively corroborate ongoing myogenic maturation and progressive structural reinforcement within the engineered construct.

**Figure 2.**
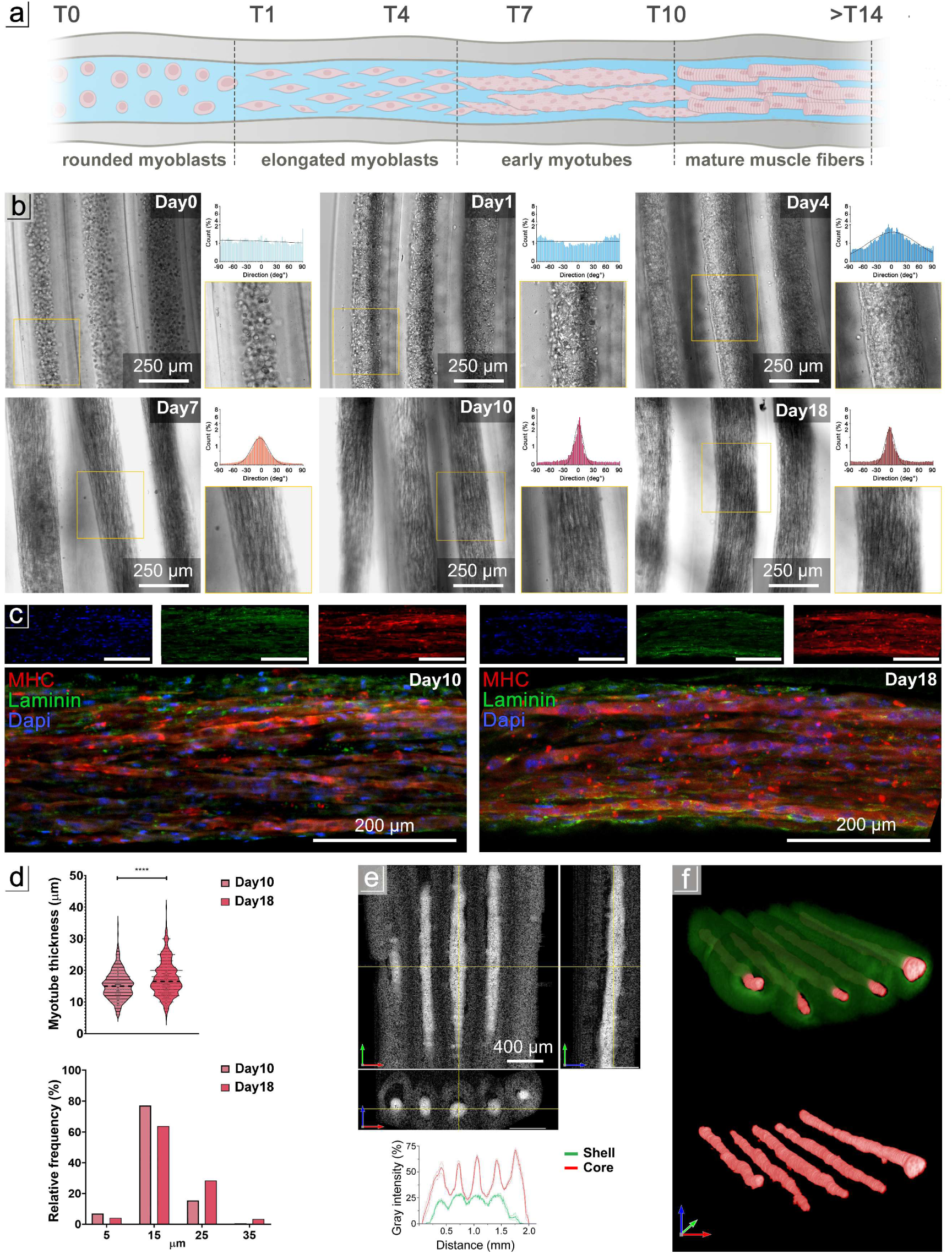
Generation and structural maturation of core-shell muscle bundles. a) Timeline of myogenic progression in core-shell fibers fabricated via microfluidic-assisted wet-spinning, from cell encapsulation (T0) to anisotropic myobundle formation (>T14); b) Brightfield microscopy images showing the transition from dispersed myogenic precursors (T0) to aligned myotubes (>T14). To the right of each brightfield image, the quantitative directionality analysis displays the angle distribution of fiber alignment for the corresponding time point; c) Immunofluorescence staining for myosin heavy chain (MHC, red), laminin (green), and nuclei (DAPI, blue) at Day 10 and Day 18, showing multinucleated myotubes and laminin-rich basement-membrane organization; d) The graphs show the quantification of myotube thickness at Day 10 and Day 18 (left; mean ± SD) and the corresponding relative thickness distribution for each time point (right). Student’s t-test was used to evaluate the differences between means; significant differences: *p < 0.05, ***p < 0.001,****p < 0.0001; e) Optical coherence tomography (OCT) images highlighting core–shell architecture, with increased signal intensity in the core region. The grayscale intensity profile across the fiber cross-section shows the spatial separation of core and shell compartments. f) Segmentation mask distinguishing the muscle-dense core (red) from the alginate-based shell (green).

Structural validation of the myotubes was performed using OCT imaging and grayscale intensity analysis (**Figure 2e**), which confirmed the precise core-shell compartmentalization of eSM. A segmentation mask (**Figure 2f**) was applied to distinguish the two distinct regions within the construct. This spatial mapping confirmed the compartmental integrity and the replication of structural anisotropy of the eSM. The high-intensity signals in the core region reflected the presence of compact, aligned muscle fibers, indicative of a homogeneous cell growth throughout the fiber core volume. OCT orthogonal cross-sections revealed a partial compaction of the myotube bundle within each fiber. This aspect aligns with other studies, where myogenic precursors encapsulated in hydrogel tend to undergo compaction during matrix remodeling and myotube formation [18]. In contrast, the shell showed lower intensity, consistent with its hydrogel-based composition, and showed no signs of cell infiltration. To further characterize the maturation and contractile functionality of the eSM constructs, we assessed sarcomeric organization, spontaneous contractility, and transcriptional dynamics of myogenic markers (**Figure 3**). Representative immunofluorescence imaging at the latest differentiation stage (Day 18) demonstrated clear α-sarcomeric actinin striation patterns, confirming the establishment of organized sarcomeres within multinucleated myotubes (**Figure 3a**). Quantification of sarcomere length revealed comparable Z-disk spacing between the two time points (Day 10 and Day 18), with values of approximately 2 µm (**Figure 3b**), consistent with physiological sarcomere dimensions and indicating that a mature contractile architecture was already established by Day 10 [19]. Line-profile fluorescence intensity analysis further supported this periodic organization, highlighting the regular alternation of α-actinin-positive Z-disks. Functionally, mature constructs exhibited spontaneous contractions (i.e., twitching). Displacement analysis performed by overlaying relaxed and contracted fiber masks revealed coordinated motion across all fibers within the bundle, highlighting synchronous activation and mechanical coupling (**Figure 3c**).

**Figure 3.**
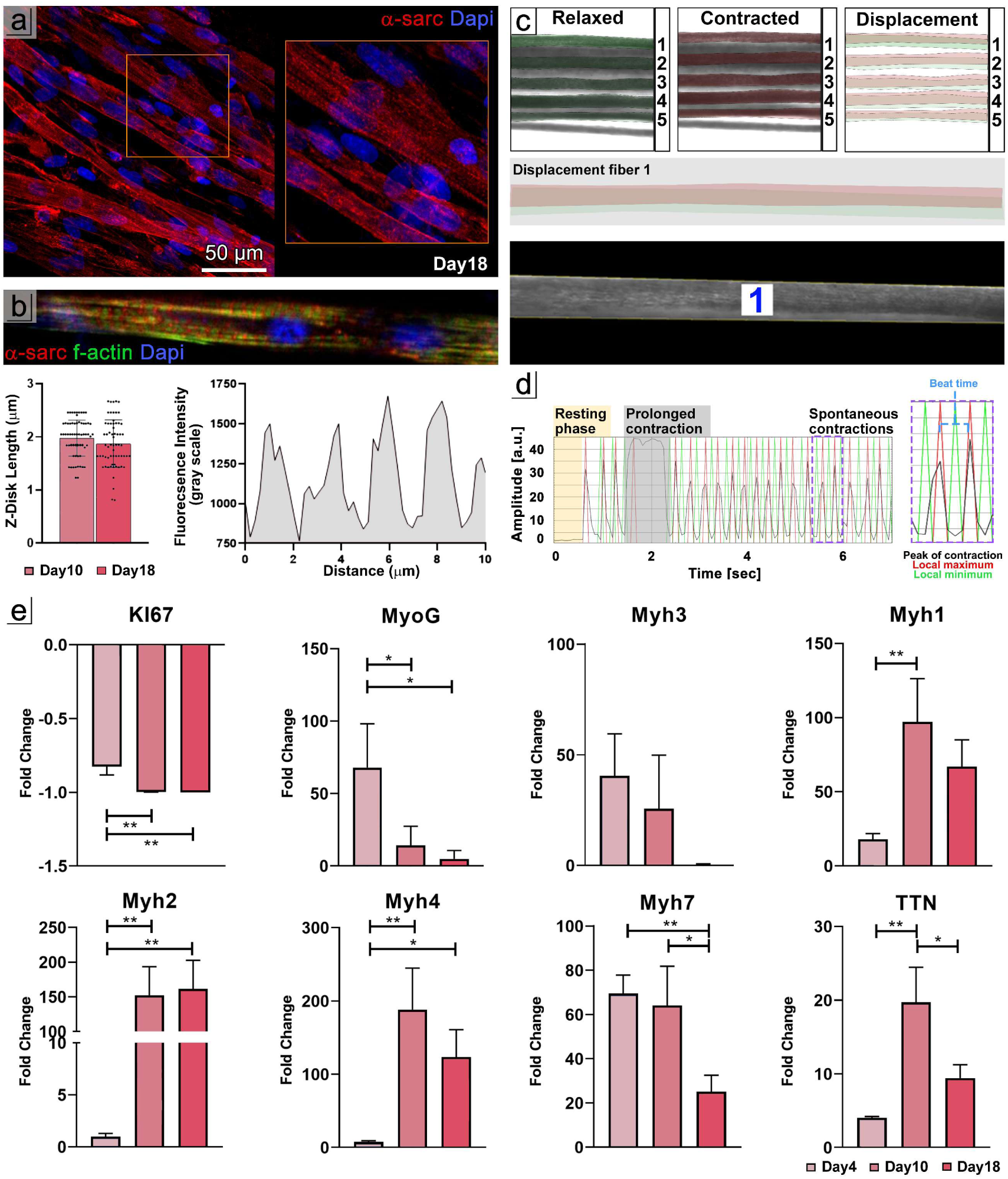
Structural, functional, and molecular characterization of eSM constructs. a) Immunofluorescence staining at Day 18 showing α-sarcomeric actinin (red) and nuclei (DAPI, blue), demonstrating organized sarcomeric structures; b) Z-disk length analysis at Day 10 and Day 18, showing mean ± SD values and representative fluorescence-profile intensity measurements illustrating Z-disk periodicity. c) Displacement analysis showing relaxed versus contracted fiber masks and displacement traces; Spatiotemporal kymograph and contraction trace generated using Myocyter for the fiber number 1; d) Relative gene expression of proliferation (Ki67) and myogenic differentiation markers (MyoG, Myh3, Myh1, Myh2, Myh4, Myh7, and TTN) at Day 4, Day 10, and Day 18, expressed as fold-change versus Day 0 (mean ± SD). One-way ANOVA was used to evaluate differences among the different experimental groups. Statistical significance was defined as *p < 0.05, ***p < 0.001,****p < 0.0001.

For clarity, a representative displacement trace from a single fiber (**fiber #1**) is shown in the main figure, while the full displacement analysis across all five fibers is reported in **Supplementary Figure 1**. Spatiotemporal kymographs further confirmed simultaneous displacement events along individual fibers. Individual fiber traces demonstrated consistent, homogeneous contraction amplitudes and preserved rhythmicity. Automated contraction profiling performed using the Myocyter ImageJ plugin (**Figure 3c**), showed a characteristic progression from an initial resting phase to spontaneous twitching events, including a prolonged contraction phase followed by repetitive cyclic contractions. Quantitative contraction metrics (**Supplementary Table 2**) revealed an average beat time of ∼ 0.5 s, a contraction frequency of ∼ 3.1 Hz, and a mean contraction amplitude of ∼ 21.5 a.u., together with a peak time of 0.161 s, a contraction time of 0.08 s, and a relaxation time of 0.124 s, values consistent with active myofiber behavior. Together, these data demonstrate that the engineered muscle fibers develop coordinated, spontaneous contractility, supporting the establishment of a functionally mature muscle architecture within the engineered constructs. Gene expression analysis corroborated these findings, revealing a temporal progression characteristic of myogenic differentiation (**Figure 3d**). Fold-change values were calculated relative to the wet-spun 3D eSM samples at day 0. The proliferation marker Ki67 was downregulated from day 4 onward, reflecting exit from the cell cycle and commitment toward differentiation [20]. The early myogenic regulator MyoG (myogenin) peaked at day 4 and decreased at later stages, consistent with its role in initiating myoblast fusion [21]. Similarly, the embryonic myosin heavy chain Myh3 was upregulated at day 4 and progressively reduced by day 18, supporting a transition toward more mature fiber identity [21]. In contrast, the expression of multiple adult myosin isoforms-including Myh1 (fast type IIx), Myh2 (fast type IIa), Myh4 (fast type IIb), and Myh7 (slow type I)-markedly increased at day 10, along with Titin (TTN), a key determinant of sarcomere assembly and mechanical stability [22].

By day 18, expression levels of adult myosins and TTN partially returned toward baseline, consistent with the establishment of a stable differentiated phenotype following a peak in terminal myogenic commitment at day 10.

Collectively, these structural, functional, and molecular readouts confirm that the engineered muscle constructs undergo efficient differentiation, establish physiologically relevant sarcomere organization by day 10, and progress toward a stable, contractile, and transcriptionally mature phenotype over time. Altogether, these results highlight the capability of our microfluidic-assisted wet-spinning platform to produce highly anisotropic skeletal muscle constructs with biomimetic architectural and functional properties in a straightforward yet effective manner.

### 3.3 Functional Microvascular Seeds production

To fabricate monodisperse hydrogel beads in large quantities, we designed a millipede microfluidic chip that operates via step emulsification. This multi-channel configuration ensures inherent stability in the emulsification process, reducing sensitivity to flow-rate fluctuations and leading to uniform droplet sizes and reliable performance. The design and operational principle of the millipede microfluidic chip are illustrated in **Figures 4a and 4b. Figure 4a** depicts a schematic rendering of the millipede layout, including a magnified view of the conical nozzle terminations at the outlet. In this configuration, a 5% w/v GelMA solution was used as the droplet phase, and the highly parallelized array of nozzles enabled efficient shearing and detachment of GelMA droplets in proximity of the step region. Once formed, the droplets were encapsulated within the oil phase and guided through a progressively widening channel toward the outlet, ensuring uniform bead formation and stable downstream transport with minimal coalescence or size variability (**Figure 4b)**. Following UV-induced crosslinking and surfactant removal, the resulting beads retained a well-defined spherical morphology and sharp contours, as observed in **Figure 4c**, indicating efficient photopolymerization.

**Figure 4.**
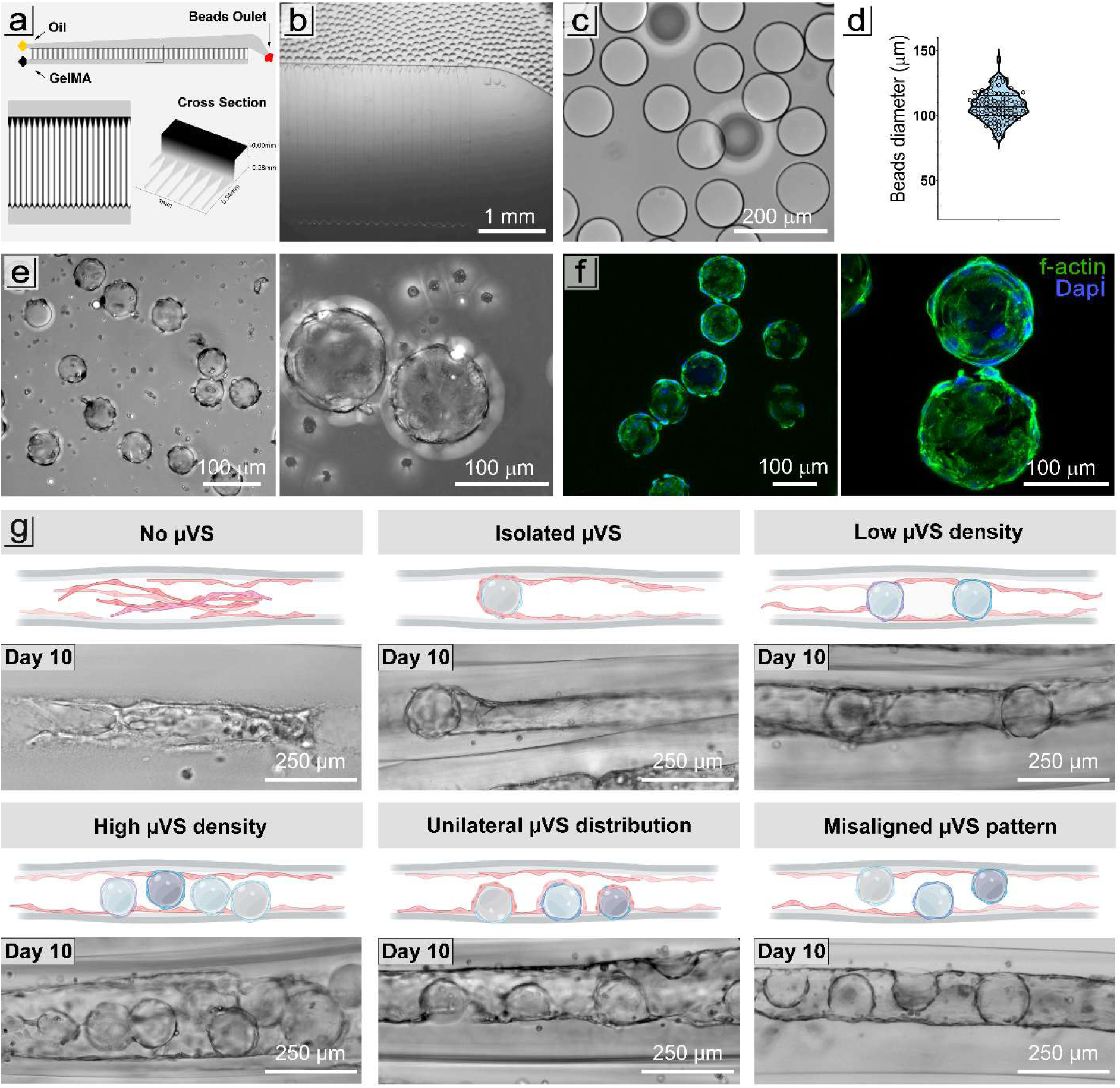
Microvascular seed production. a) Schematic rendering of the millipede microfluidic chip with zoomed-in view of conical microchannels geometry for uniform droplet formation and a cross-sectional view of the chip structure; b) Brightfield image showing synchronized GelMA droplet generation; c) Crosslinked GelMA beads after surfactant removal; d) Box plot showing bead diameters distribution (mean ∼110 µm); e) Brightfield images of microvascular seeds 4 h after endothelial cell seeding, shown at low and high magnification; f) Confocal images of the same microvascular seeds stained for F-actin (green) and DAPI (blue); g) representative distribution patterns of µVS within wet-spun fibers at day 10 illustrating six stochastic spatial configurations: (i) no µVS, (ii) isolated µVS, (iii) low-density, (iv) high-density, (v) unilateral distribution, and (vi) misaligned arrangement. The renderings were created with Biorender.com.

Quantitative analysis of bead diameter distribution (**Figure 4d**) revealed a mean size of approximately 110 µm, with a standard deviation of 11 µm. These beads were generated using flow rates of 100 µL/min for both the GelMA solution and the oil phase, reaching a throughput of approximately. 1.4 ×10^5^ beads/min. These flow rates and bead dimensions were selected following a series of preliminary optimization experiments (data not shown), in which multiple parameter combinations were tested to identify conditions that ensured maximal monodispersity, stability, high-throughput performance, and an optimal bead size for efficient cell adhesion. The resulting monodispersity ensured a consistent surface-area-to-volume ratio, which is crucial for achieving homogeneous cell seeding and predictable behavior in downstream endothelialisation processes. To generate microvascular seeds (µVS), endothelial cells (HUVECs) were dynamically seeded onto the surface of crosslinked GelMA beads and cultured for 4 hours. Brightfield microscopy at both low and high magnification (**Figure 4e**) confirmed the preservation of bead morphology after seeding, demonstrating the proper structural integrity essential for subsequent handling and incorporation of microvascular seeds into more complex tissue constructs. Confocal immunofluorescence analysis (**Figure 4f**), performed after an overnight incubation following cell seeding, revealed a uniform endothelial coverage across the bead surface, with F-actin staining (green) highlighting organized cortical cytoskeletal structures and DAPI staining (blue) showing evenly distributed nuclei. This finding indicated efficient cell adhesion and early cytoskeletal remodeling on GelMA beads. Notably, during preliminary optimization experiments, we observed that HUVECs did not adhere to GelMA beads with diameters of 70–80 µm or less. This behavior may be attributed to the increased surface curvature of smaller beads, which likely limits cells’ ability to spread and reorganize their cytoskeleton on highly curved substrates. The microscale curvature of the selected beads (110 µm in diameter) appeared well-suited to support endothelial cell spreading and anchorage, providing a favorable topographical interface for the formation of a stable cellular monolayer resembling an endothelium, which in turn supports their intended use as modular, endothelialized building blocks in vascularized tissue constructs. The results also show that the millipede microfluidic chip enables high-throughput and reproducible manufacturing of size-controlled GelMA beads, coupled with efficient HUVEC seeding and full microbead functionalization, ultimately positioning the resulting µVS as optimal building blocks for advanced biofabrication workflows.

To test our hypothesis that the µVS can assemble into functional vasculature, the pre-endothelialized seeds were incorporated into the pro-angiogenic core bioink and subsequently processed on the RoWS bioprinting platform. Building on this strategy, µVSs were encapsulated at 50.000 seeds/mL concentration within a fibrin-based bioink together with resuspended HUVECs (10^7^ cells/mL) and extruded as the core phase. The concentration of microvascular seeds was optimized to ensure homogeneous dispersion while preventing aggregation or nozzle clogging. As one can easily infer, the spatial distribution of µVS within the core cannot be actively controlled, as they are resuspended during the preparation of the pro-angiogenic bioink and subsequently processed via wet spinning. Consequently, we performed a preliminary qualitative analysis to investigate how this stochastic arrangement, present in each sample, affects locally vascular morphogenesis, providing an overview of the diverse structural outcomes observed. **Figure 4g** summarizes six representative scenarios at day 10 of culture. In the no µVS condition, endothelial cells lacked anchoring sites and formed a disordered, branched network with no clear signs of lumen formation along the core [3]. In the isolated µVS case, the µVS served as a local anchoring point for endothelial organization, promoting the formation of a lumen segment that extended unidirectionally from the seed, with clear longitudinal orientation. At low density, adjacent µVS were bridged by a continuous endothelial lining, producing a homogeneous, well-defined lumen that connected successive seeds and maintained a uniform caliber along the fiber. As µVS density increased, spatial crowding led to an expansion of the core compartment. Despite this increase in seed number, the vascular structure remained well-organized, with endothelialisation incorporating multiple µVS within the lumen to form a coherent, fully connected vascular channel. At intermediate µVS densities, two distinct spatial configurations were commonly observed. When µVS were uniformly aligned along one core–shell interface (unilateral distribution), lumen formation remained uninterrupted and of relatively uniform diameter. The endothelium tended to position the µVS to one side, growing over part of their surface while preserving an overall circular architecture. Conversely, when µVS were misaligned and alternately positioned on opposite interfaces, the endothelial monolayer still formed a continuous lumen but displayed local curvature and mild tortuosity, reflecting regions where µVS were variably excluded from or incorporated into the vascular channel.

Collectively, these observations demonstrate that µVS spacing and positioning act as effective geometric and adhesive cues that guide endothelial self-organization. Regardless of the specific configuration, lumenization consistently occurred, yielding continuous and morphologically coherent vascular structures. Low-to-intermediate µVS densities favored uniform, well-defined lumens, whereas higher densities or misalignment modulated lumen size and curvature without compromising overall vascular integrity or functionality.

### 3.4 Biofabrication of tubular microvessel-mimicking model

After the preliminary tests, we proceeded with a more thorough analysis of vasculature structures obtained with µVS. The schematic rendering in **Figure 5a** illustrates the sequential progression of vascular morphogenesis orchestrated by the µVS within the engineered constructs. Following extrusion, µVS and single HUVECs are stochastically distributed within the fibrin-based core (T0). As culture progresses, the system undergoes a dynamic self-organization process reminiscent of early vasculogenic events. By day 3 (T3), endothelial cells initiate directional migration and sprouting from the µVS surface, forming nascent multicellular extensions that bridge adjacent seeds and establish the first intercellular connections. By day 6 (T6), these extensions develop into cohesive, aligned assemblies exhibiting early signs of lumen initiation and spatial organization. At day 12 (T12), the endothelial networks transition into continuous, lumenized tubular architectures extending longitudinally along the core, indicative of mature microvessel-like organization and progressive stabilization of the endothelial compartment. These dynamic transitions are shown in **Figure 5b**, where brightfield images illustrate the progression from dispersed cell-seed aggregates to organized, tubular endothelial structures. Confocal immunofluorescence analysis substantiated these morphological observations (**Figure 5c**). The simultaneous expression of CD31 (red) and von Willebrand Factor (vWF, green) confirmed endothelial identity and progressive junctional maturation. By day 6 (T6), broad and continuous CD31-positive regions were already evident, delineating extensive areas of interconnected endothelium and suggesting the establishment of stable cell-cell junctions across the forming vessel wall. By day 12 (T12), CD31 staining became uniformly distributed along the whole lumen circumference, defining a coherent and mature endothelial monolayer [23,24]. The concurrent expression of vWF, reflected enhanced secretory activity and the acquisition of a functionally quiescent endothelial phenotype, indicative of lumen stabilization and vascular maturation. To further resolve the internal organization of these engineered microvessels, optical coherence tomography (OCT) imaging was employed (**Figure 5d**). Orthogonal cross-sections revealed hollow, circular lumens extending uninterruptedly along the fiber length. The sharp contrast between the lumen and surrounding matrix confirmed the structural coherence and the absence of collapse, occlusion, or branching, hallmarks of controlled endothelial morphogenesis. Collectively, these results demonstrate that the µVS acted as geometric and biochemical nucleation foci, guiding the self-assembly of endothelial cells into hierarchically ordered, perfusion-relevant microvascular structures.

**Figure 5.**
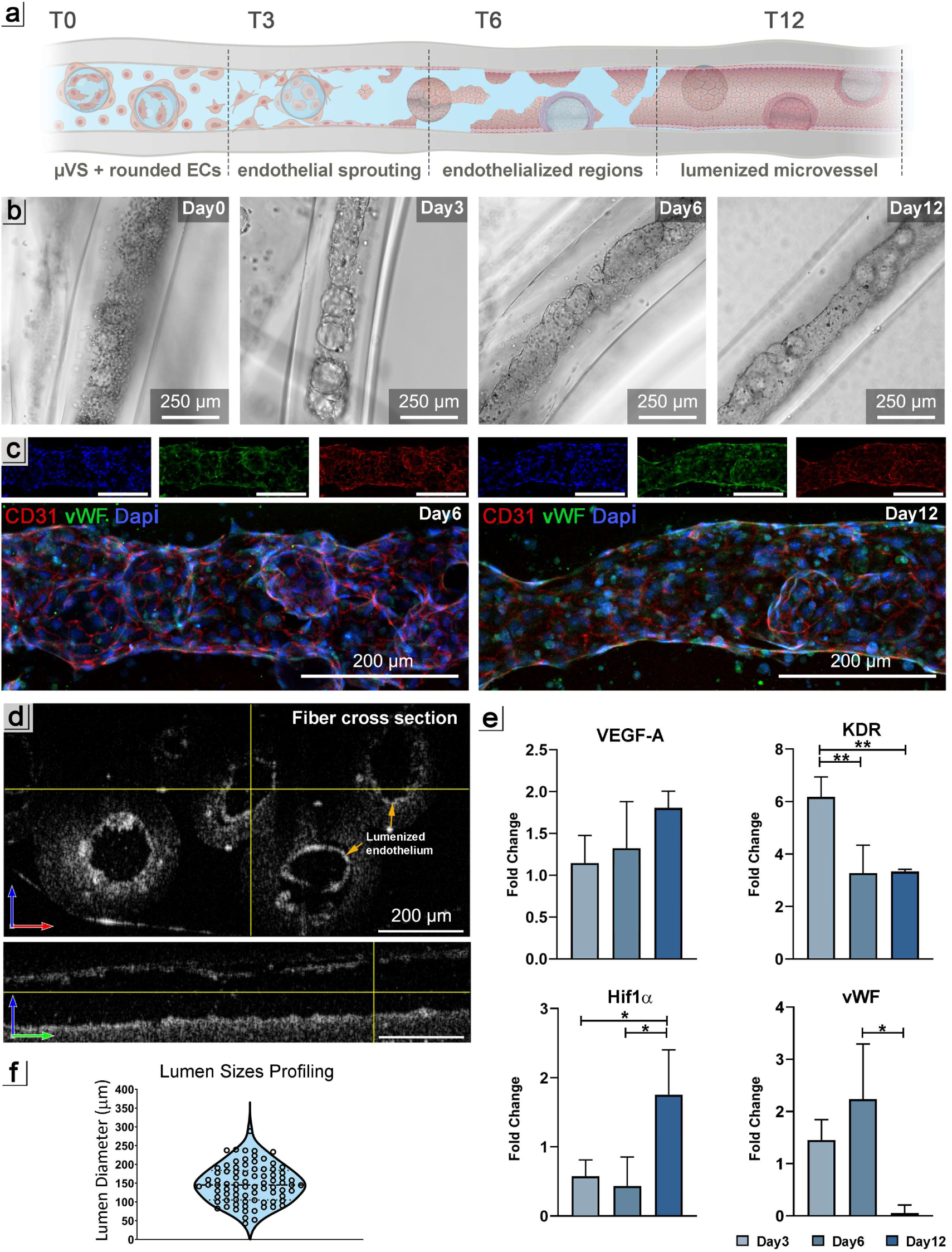
Biofabricated tubular microvessels. a) Schematic timeline rendering illustrating the temporal progression of endothelial morphogenesis within core-shell constructs, from initial dispersion (T0) to the formation of continuous lumenized networks (T12). b) Brightfield micrographs documenting the morphological evolution of endothelial structures: at T0, dispersed µVS and HUVECs; at T3, early multicellular extensions bridging adjacent µVS; at T6, extensive regions of continuous endothelium; and at T12, fully organized, lumenized microvessel-like architectures; c) Confocal immunofluorescence staining for CD31 (red), vWF (green), and nuclei (DAPI, blue) showing progressive endothelial maturation. By T6, broad CD31-positive regions delineate interconnected endothelial domains, while by T12, CD31 expression becomes circumferential and homogeneous, defining a mature, continuous endothelial monolayer with uniform vWF distribution; d) Optical coherence tomography (OCT) images providing transverse and longitudinal views along the fiber axis, revealing hollow, circular lumens distributed throughout the construct and aligned with the extrusion direction.; e) Lumen diameter distribution of the engineered microvessels at T12, showing a consistent range of 100–200 µm, compatible with physiological arteriole and venule dimensions; f) Relative gene expression of angiogenic and endothelial markers (VEGF-A, KDR, Hif1α, vWF) at day3, 6, 12 of culture, illustrating the transcriptional progression from early angiogenic activation to stabilized endothelial phenotype (mean ± SD). A one-way ANOVA was used to evaluate differences among the experimental groups. Statistical significance was defined as *p < 0.05, ***p < 0.001,****p < 0.0001.

To quantitatively assess the structural maturation of the engineered microvessels, lumen diameter profiling was performed (**Figure 5f**). The analysis revealed a narrow and consistent distribution centered around 100-200 µm, values that fall within the physiological range of arterioles and venules. This dimensional fidelity is particularly relevant when compared to conventional endothelial self-assembly approaches, which typically yield capillary-like networks with diameters of less than 10 µm [25]. Such microvasculatures, though morphologically intricate, often lack structural robustness and exhibit limited perfusability, thus constraining their capacity to sustain thick, metabolically active tissues [25]. As highlighted by Paradiso et al. [3] microvascular architectures with diameters below the diffusion threshold are insufficient to maintain high viability in engineered thick constructs, where rapid perfusion is essential to ensure effective oxygen and nutrient delivery and waste removal. In contrast, the lumenized constructs generated through our µVS-guided strategy exhibited stable, pre-patterned microchannels of physiologically relevant size, providing a structural framework amenable to subsequent perfusion and potential integration with host vasculature upon implantation. These features collectively highlight the advantage of incorporating pre-endothelialized µVS within the wet-spinning process, enabling the formation of functionally scalable vascular units rather than fragile capillary beds. To further characterize the endothelial phenotype and verify the transcriptional progression toward vascular maturation, quantitative RT–PCR was performed on constructs harvested at days 3, 6, and 12 (**Figure 5e**), with gene expression levels reported as fold changes relative to day 0. Vascular Endothelial Growth Factor A (VEGF-A) expression showed a progressive, time-dependent increase over the culture period, reaching nearly a 2-fold increase at day 12 compared to baseline. This upregulation is consistent with the established role of VEGF-A in supporting endothelial survival, vascular remodeling, and long-term maintenance of endothelial function, beyond its initial involvement in angiogenic sprouting [26,27]. A more pronounced and dynamic regulation was observed for Kinase Insert Domain Receptor (KDR/VEGFR2), which showed a marked early upregulation, with an approximately 6-fold increase detectable by day 3. This early peak was followed by significant attenuation at days 6 and 12 compared to day 3, while remaining above day 0 levels. Such a transient expression profile is characteristic of an early endothelial activation phase, associated with heightened VEGF responsiveness, which is progressively reduced as endothelial cells transition toward a more stable, mature phenotype [28,29]. In contrast, Hypoxia-Inducible Factor 1 Alpha (HIF1α) expression remained comparable to baseline at day 3 and day 6 but showed a pronounced and significant upregulation at day 12 relative to both earlier time points. Notably, HIF1α activation is not exclusively driven by hypoxic stimuli, as it can also be induced by growth factors and by signaling pathways, including VEGF-dependent pathways, in endothelial cells. In this context, the delayed HIF1α upregulation observed here may reflect VEGF-driven transcriptional reinforcement mechanisms operating during prolonged culture, consistent with previous reports demonstrating that VEGF signaling can induce HIF1α expression and activity independently of hypoxia, thereby sustaining angiogenic and endothelial adaptive programs [30]. Finally, von Willebrand Factor (vWF) displayed a distinct temporal pattern, with a peak of expression at day 6 followed by a return to baseline levels at day 12. Given that vWF is commonly associated with endothelial activation and early vascular organization, its transient upregulation followed by normalization indicates a shift from an activated endothelial state toward a more stabilized, functionally reorganized endothelium [31]. Importantly, beyond its role as an endothelial marker, vWF has been implicated as a regulator of angiogenesis and vascular stabilization, with reduced vWF levels being associated with excessive yet potentially dysfunctional angiogenic responses; therefore, the observed normalization at day 12 is consistent with the progression toward a more controlled and stabilized endothelial phenotype [31,32]. Altogether, these results highlight the µVS-guided wet-spinning strategy as an effective biofabrication approach for generating hierarchically organized, lumenized microvascular structures, in which spatial patterning, morphological stabilization, and transcriptional maturation converge to achieve a functional endothelial phenotype.

### 3.5 Integration of vascular and muscle-mimicking compartments

A critical milestone in the development of biomimetic muscle constructs is the integration of vascular and muscular compartments into a unified, hierarchically organized tissue. The workflow for generating vascularized skeletal muscle constructs using the 3D RoWS platform is schematically represented in **Figure 6a**. As previously described, skeletal muscle fibers were fabricated using a myogenic bioink and cultured for 18 days in a muscle-specific medium to first promote myoblast proliferation, followed by differentiation and maturation into multinucleated myotubes exhibiting key structural and molecular features of native skeletal muscle. In parallel, µVS, together with single HUVECs, were incorporated into a pro-angiogenic bioink and cultured for 12 days, resulting in the formation of lumenized microvascular structures lined by a continuous endothelial monolayer. Demonstrating the preservation of cell viability and structural integrity in each compartment under co-culture conditions is essential. Indeed, preserving these biological features over time is crucial for bridging the *in vitro* phase with potential *in vivo* implantation in animal or human models. For this purpose, eSM constructs at day 18 of maturation and tubular microvessel constructs at day 12 were assembled into a single construct, transferred to the co-culture medium, and maintained for an additional 7 days to assess cell viability, structural stability, and maintenance of compartment-specific organization under shared culture conditions. As shown in **Figure 6b**, both eSM and tubular microvessels exhibited high cell viability, with live-cell percentages exceeding 80%, while maintaining their structural integrity and anisotropy. These observations indirectly suggest that tubular microvessel structures can sustain metabolic activity and maintain lumen robustness under culture conditions simulating myogenic differentiation, and that, conversely, the presence of pro-angiogenic factors does not impair muscle differentiation. As shown in **Figure 6c**, bright-field microscopy reveals the coexistence of aligned myofibers and adjacent vessel-like structures within the same construct, providing clear visualization of their parallel spatial distribution. Importantly, the ring-shaped configuration of the constructs, composed of concentrically organized bundles of aligned fibers, inherently facilitates their modular combination into larger composite structures while preserving anisotropy and spatial definition. Consequently, this spatial organization recapitulates the native skeletal muscle environment, in which microvessels are positioned alongside contractile fibers to meet metabolic and functional demands [33]. To further characterize the spatial organization and phenotypic identity of the two compartments, immunofluorescence analysis was performed using laminin (green), myosin heavy chain (MHC, red), and CD31 (magenta) (**Figure 6d**). Confocal images confirm the effective assembly of the vascularized muscle construct, showing MHC-positive, well-aligned myofibers and distinct endothelial structures, with both compartments clearly identifiable and structurally preserved following integration. Quantitative RT-PCR analysis (**Figure 6e**) was performed following the co-culture period to confirm and monitor the transcriptional changes induced by endothelial–muscle interaction. For the microvascular component, a modest increase in VEGF-A and HIF1α expression was observed in co-cultured constructs compared to monoculture controls, although the magnitude of this change remained below 1-fold and did not indicate strong transcriptional activation. In contrast, KDR (VEGFR2) and, more prominently, vWF showed a clear upregulation in the co-culture condition. From a biological perspective, the selective upregulation of KDR and vWF, rather than a broad activation of pro-angiogenic genes, is consistent with a shift toward endothelial stabilization and functional organization rather than active sprouting. KDR upregulation may reflect increased endothelial cell sensitivity to VEGF signaling cues from muscle tissue, even in the absence of strong VEGF-A induction. Notably, vWF upregulation is indicative of endothelial maturation and vascular homeostasis, as previously described. Its increased expression in co-culture suggests that endothelial cells adopt a more functionally organized and stabilized phenotype in the presence of differentiated muscle tissue, rather than an activated or proliferative angiogenic state. In the muscle compartment, expression of late structural and contractile markers, including Myh1, Myh4, and TTN, showed mild downregulation in co-cultured constructs compared with monoculture controls, whereas Myh2 and Myh7 showed a slight increase. However, these variations remained limited in magnitude and did not reach statistical significance. This transcriptional profile suggests that muscle identity and differentiation status were largely preserved during co-culture, with only subtle remodeling at the gene expression level, consistent with the use of an already mature and differentiated muscle compartment, for which no abrupt transcriptional changes would be expected over short co-culture periods. The modest increase in Myh2 and Myh7, which are associated with more oxidative and slow/intermediate fiber phenotypes, may therefore reflect a mild adaptive adjustment to the presence of a vascular compartment, rather than a shift in muscle differentiation state, potentially mirroring early aspects of metabolic or functional coupling between muscle fibers and endothelial cells.

**Figure 6.**
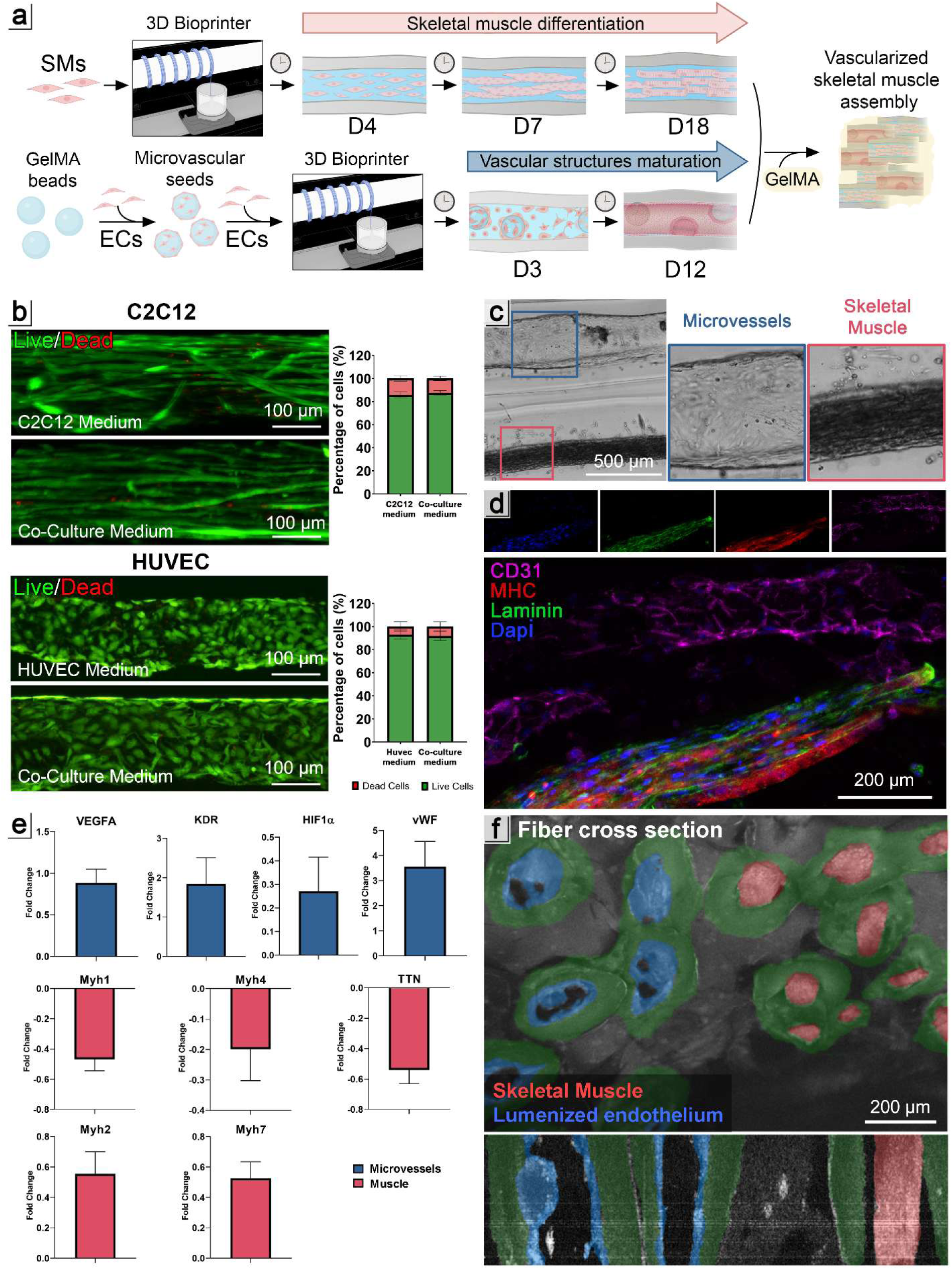
Vascularized skeletal muscle construct assembly. a) Schematic representation of the biofabrication workflow, illustrating the separate generation of myogenic and microvascular components followed by their assembly into a vascularized skeletal muscle construct; b) Live/Dead assay performed on both muscle and vascular constructs after an additional 7 days of culture following their respective maturation phases, maintained in their specific media (myogenic or endothelial) and in the co-culture medium. Quantitative analysis indicates cell viability exceeding 80 % in both compartments after the extended culture period under all conditions, confirming structural preservation and metabolic stability; c) Brightfield image showing the integration of pre-matured muscle fibers (day 18) and vascular structures (day 12) into a single construct; d) Immunofluorescence staining of the vascularized muscle construct showing laminin (green), myosin heavy chain (MHC, red), CD31 (magenta), and nuclei (DAPI, blue); e) Gene expression of endothelial (VEGF-A, KDR, HIF1α, vWF) and myogenic markers (Myh1, Myh2, Myh4, Myh7, TTN) in co-cultured constructs after 7 days of co-culture, expressed relative to corresponding monoculture constructs. Data are shown as fold change (mean ± SD), significant differences: *p < 0.05, **p < 0.01, ***p < 0.001; f) OCT imaging with applied segmentation mask showing cross-sectional and longitudinal views of the construct. The shell portion is highlighted in green, muscle fibers are visible in red, and the endothelial monolayer is visualized in blue.

Overall, the gene expression data indicate that short-term co-culture does not induce major transcriptional reprogramming in either compartment, but rather promotes selective, compartment-specific adjustments. Finally, OCT imaging (**Figure 6f**) was employed to obtain a comprehensive 3D visualization of the assembled construct, enabling simultaneous assessment of its overall morphology and internal architecture. The applied segmentation mask enabled the clear identification of both muscle fibers (red) and lumenized vascular structures (blue), which were spatially compartmentalized by the shell region (green). This analysis confirmed the effectiveness of the assembly strategy in preserving spatial separation and structural integrity of the individual compartments within the single unified construct. The resulting configuration, featuring aligned myofibers and tubular endothelial structures in parallel orientation, recapitulates key morphological and biological features of native skeletal muscle. These findings contrast with previous co-culture models, in which joint environments limited maturation efficiency. Importantly, our platform also circumvents key limitations of conventional 3D bioprinting approaches to fabricate heterogeneous tissue interfaces, such as insufficient resolution for sub-millimeter vascular features, multi-nozzle complexity, and long fabrication times [34]. The microfluidics-assisted wet-spinning system offers a scalable and reproducible method for generating anisotropic, spatially defined tissues, suitable for regenerative therapy applications [18–20], such as the reconstruction of vascularized tissues in volumetric muscle loss, as well as for modeling physiologically relevant, complex muscle microenvironments *in vitro*.

## 4. CONCLUSIONS

In this study, we introduced a modular biofabrication strategy that enables the independent maturation and subsequent integration of muscle and vascular compartments into a unified, hierarchically organized construct. Using the RoWS platform, we fabricated precise core-shell architectures supporting: i) the differentiation of aligned, multinucleated myotubes displaying organized sarcomeres, spontaneous coordinated contractions, and a transcriptional signature consistent with terminal myogenic maturation; ii) the establishment of µVS as modular, pre-endothelialized units that act as ready-to-use vascular modules, potentially compatible with multiple biofabrication strategies, accelerating the assembly of endothelialized architectures; iii) the formation of µVS-guided, lumenized microvessel structures exhibiting physiologically relevant calibers (≈100-200 µm), continuous endothelial monolayers, and gene expression associated with vascular stabilization. Importantly, both the muscle and microvascular compartments maintained high viability, spatial organization, and their established phenotypic identities after one week of co-culture, with only subtle, compartment-specific transcriptional adjustments. This demonstrates the compatibility of this biologically heterogeneous system under shared culture conditions while maintaining the maturation state of each component. All together, these findings highlighted how decoupled maturation followed by spatially controlled assembly overcomes the limitations of conventional co-culture systems, generating a scalable and reproducible method for engineering anisotropic, vascularized muscle tissues. This model could serve as a supporting tool for regenerative strategies in the context of VML, while providing a physiologically relevant *in vitro* platform to study endothelial–muscle crosstalk, assess angiogenic and myogenic cues, and evaluate candidate therapeutics in a controlled 3D microenvironment.

It is important to acknowledge that the GelMA-based beads forming the core of the µVS may initially act as partial lumen obstructions. Within the experimental timeframe, in fact, no substantial degradation was observed under standard *in vitro* conditions. Future studies will therefore explore alternative biomaterials with enhanced hydrolytic or enzymatic degradability to promote more efficient *in vitro* remodeling. Conversely, GelMA degradation *in vivo* has been investigated in various studies, which demonstrated a resorption time ranging from 2 to 4 weeks [35–37] depending on hydrogel formulation, enzymatic activity, inflammatory signaling, and immune cell infiltration at the implantation site. These considerations suggest that, in a physiological *in vivo* setting, GelMA-based µVS may undergo rapid remodeling and clearance, supporting the feasibility of future *in vivo* applications of lumenized microvessels generated through µVS-guided vascular assembly. Finally, owing to their micrometre-scale dimensions and structural stability, µVS may also serve as modular vascular bioadditives for bioink formulations, enabling pre-endothelialized building blocks to be integrated with diverse biofabrication platforms. Such an approach could expand their applicability beyond the present system and further enhance their translational potential within the biofabrication field. Overall, this work demonstrates that integrating pre-vascularized and pre-matured muscle modules through microfluidic-assisted wet-spinning enables the fabrication of structurally biomimetic, functionally mature tissues, narrowing the gap between engineered *in vitro* models and clinically relevant biofabricated tissues.

## Supporting information

Three-dimensional reconstruction of a microvascular segment at day 12 showing endothelial monolayer organization, including external and luminal views

Spontaneous contractile activity of rotary wet-spun skeletal muscle constructs under differentiation conditions at day 18.

## 5. ACKNOWLEDGEMENTS

This work was supported by the National Science Centre Poland (NCN) within SONATA BIS 12 (project no. 2022/46/E/ST8/00284 to M.C.). Figures 4 and 5 were partly created with BioRender.com.

## Notes

### Competing Interest Statement

The authors have declared no competing interest.

