## Supplementary Information for "Modular biofabrication of a vascularized skeletal muscle model through endothelialized microvascular seeds"

**Supplementary Table 1**. Murine and human primer sequences for qRT-PCR analysis.

| Gene | Forward primer | Reverse primer |
|---|---|---|
| murine Tbp | CAGAACAACAGCCTTCCACC | GCTGCTGTCTTTGTTGCTCT |
| murine Ki67 | CATGAGGATGGAAGCAAGCC | TGCTGTTCTACATGCCCTGA |
| murine MyoG | CGCAGGCTCAAGAAAGTGAA | GTGGGAGTTGCATTCACTGG |
| murine Myh1 | GACTACAACATCGCTGGCTG | CTTGGCCCCTTTCTTTCCAC |
| murine Myh2 | AACATCTGTCTTTGTGGCGG | CTTGTCGTACTTGGGAGGGT |
| murine Myh3 | GCAAGTCGGAAAAGGAGAGG | GGTTCATGGCATACACGTCC |
| murine Myh4 | AAGACTGTGAACACGAAGCG | TTCCTCACAGTCTTGGCGTT |
| murine Myh7 | TGTGCCCGATGACAAAGAAG | GCCATGTCCTCGATCTTGTC |
| murine Ttn | GATCATTGTCCCTGCGTCAC | TCATTTCGAGCCTGGAACCT |
| human GUS | GGAATTTTGCCGATTTCATGA | CCGAGTGAAGATCCCCTTTTT |
| human VEGF-A | TTGCCTTGCTGCTCTACCTCCA | GATGGCAGTAGCTGCGCTGATA |
| human KDR | GGAACCTCACTATCCGCAGAGT | CCAAGTTCGTCTTTTCCTGGGC |
| human HIF1α | TATGAGCCAGAAGAACTTTTAGGC | CACCTCTTTTGGCAAGCATCCTG |
| human vWF | CCTTGAATCCCAGTGACCCTGA | GGTTCCGAGATGTCCTCCACAT |

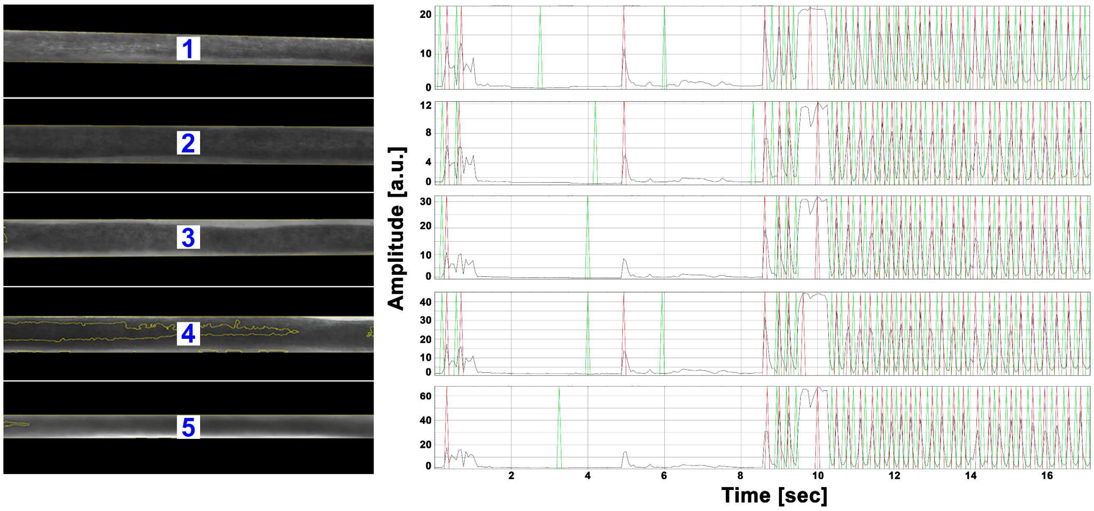


**Supplementary Figure 1. Spontaneous contraction traces of individual engineered muscle fibers.** Myocyter-derived contraction profiles for five analyzed fibers, showing amplitude (a.u.) over time (s) and highlighting consistent spontaneous rhythmic contractions across the bundle.

**Supplementary Table 2.** Quantitative contraction metrics extracted from Myocyter-based analysis of engineered skeletal muscle fibers.

| | Min | Max | Mean | StdDev |
|---|---|---|---|---|
| Beattimes [sec] | 0.250 | 5.900 | 0.581 | 1.170 |
| Frequency [1/sec] | 0.188 | 4.000 | 3.150 | 0.958 |
| Amplitudes [a.u.] | 12.666 | 32.520 | 21.599 | 3.903 |
| Peaktimes [sec] | 0.063 | 0.838 | 0.161 | 0.141 |
| Contraction [sec] | 0.050 | 0.388 | 0.087 | 0.086 |
| Relaxation [sec] | 0.063 | 0.450 | 0.124 | 0.079 |

**Supplementary Movie 1.** Spontaneous contractile activity of rotary wet-spun skeletal muscle constructs under differentiation conditions at day 18.

**Supplementary Movie 2.** Three-dimensional reconstruction of a microvascular segment at day 12 showing endothelial monolayer organization, including external and luminal views.
